# A mouse model of ATRX deficiency with cognitive deficits and autistic traits

**DOI:** 10.1101/2023.07.19.549759

**Authors:** Katherine M. Quesnel, Nicole Martin-Kenny, Nathalie G. Bérubé

## Abstract

ATRX is an ATP-dependent chromatin remodeling protein with essential roles in safeguarding genome integrity and modulating gene expression. Deficiencies in this protein cause ATR-X syndrome, a condition characterized by intellectual disability and an array of developmental abnormalities, including features of autism. Previous studies demonstrated that deleting ATRX in mouse forebrain excitatory neurons postnatally resulted in male-specific memory deficits. Here, we introduce a new model where ATRX is deleted at earlier embryonic stages, resulting in a broader spectrum of impairments, including contextual fear memory deficits, decreased anxiety, hyperactivity, as well as self-injurious and stereotyped behaviours. Sex-specific alterations were also observed, with males displaying heightened aggression and impaired sensory gating, while females exhibit social avoidance. Collectively, the findings indicate that early developmental abnormalities arising from ATRX deficiency in neurons contribute to of the presentation of autistic-like behaviours.

**Summary Statement:** Mice with embryonic loss of ATRX in excitatory neurons represent a clinically relevant model to study sexually dimorphic alterations in cognitive and autistic traits.

## INTRODUCTION

Intellectual disability (ID), defined as early onset deficits in cognitive function that lead to difficulties in daily tasks and social skills (Fahrner and Bjornsson, 2014), impacts 1-2% of school aged children (Maulik et al., 2011) and poses both social and medical challenges. Autism spectrum disorder (ASD) is another neurodevelopmental condition with high prevalence (1-2%) and is characterized as social communication difficulties with the presence of stereotyped or repetitive behaviours (Abrahams and Geschwind, 2008; Baio, J, Wiggins, L, Christensen, 2014; Bougeard et al., 2021). Many patients with neurodevelopment disorders have co-occurring conditions and often disorders overlap in their defining characteristics. Clinical studies suggest that ∼70% patients with ASD display a form of learning or intellectual disability. Additionally, anxiety and hyperactivity are associated features of ASD and 68% of patients with ASD have been reported to display aggression at some point (Bougeard et al., 2021; Fitzpatrick et al., 2016; Kas et al., 2014). These overlapping features in neurodevelopmental disorders suggest that similar molecular pathways and neural circuitry are disrupted across disorders.

Chromatin structure modulation and epigenetic mechanisms have been implicated in cognitive processes and many genes mutated in ID and ASD encode chromatin regulatory factors (Barrett et al., 2011; Feng et al., 2010; Levenson et al., 2006). Many of these chromatin remodelers are known to interact with one another in processes important for neurodevelopment and plasticity (Kochinke et al., 2016). Alpha thalassemia mental retardation, X-linked (ATRX) is an ATP-dependent chromatin remodeling protein that maintains chromatin structure integrity and regulates gene expression. Hypomorphic mutations in the *ATRX* gene result in ATR-X syndrome, a congenital syndrome that primarily affects males, as females are protected by skewed X-chromosome inactivation (Gibbons et al., 2008). Patients with ATR-X syndrome show a variety of symptoms including mild-to-severe intellectual disability, seizures, microcephaly, and dysmyelination.

In addition to intellectual disability, a subset of ATR-X syndrome patients also display autistic features (Garrick et al., 2004; Gibbons and Higgs, 2000; Gibbons et al., 1995). Moreover, *ATRX* mutations have been observed in non-syndromic patients with ID (Grozeva et al., 2015) and have been identified in patients with ASD (Gong et al., 2008; Li et al., 2017). The SFARI gene database ranks ATRX as a Category 1 (High Confidence) risk gene implicated in ASD (https://gene.sfari.org/). Based on these reports we can confidently infer that mutations in *ATRX* are highly associated with both ID and ASD.

ATRX is localized at heterochromatic regions of the genome and is involved in maintaining repressive states required for genomic integrity (Bérubé et al., 2000; Law et al., 2010; Meyer-Nava et al., 2020; Wong et al., 2010). The role of ATRX in preserving genomic integrity is crucial in actively dividing cells. Conventional knockout mice cannot be generated since ATRX-null embryonic stem cells are too unstable. Deletion of ATRX in early stages of embryogenesis or in neuronal progenitor cells results in embryonic or early post-natal death (Berube et al., 2005; Garrick et al., 2006; Seah et al., 2008). Further investigation indicated an increase in DNA damage and TP53-induced cell death upon loss of *Atrx* in neural progenitor cells (Berube et al., 2005; Seah et al., 2008; Watson et al., 2013). Therefore, to study how ATRX dysfunction leads to the clinical phenotypes observed, conditional inactivation approaches were applied to inactivate ATRX specifically in non-proliferating (post-mitotic) cells to avoid DNA replication stress-mediated cell death. We previously reported that deletion of ATRX in excitatory forebrain neurons using the αCaMKII-Cre line of mice led to male-specific deficits in long-term hippocampal-dependent spatial memory and transcriptional changes relating to synaptic regulation (Tamming et al., 2020). However, autistic phenotypes were not present in either male or female *Atrx*^CaMKIICre^ mice (Martin-Kenny & Bérubé, 2020).

This led us to hypothesize that perhaps an earlier deletion of *Atrx* might be required to replicate clinically observed cognitive deficits and features of ASD. In the current study, we utilized the NEX-Cre driver line of mice to delete *Atrx* in postmitotic neurons of the embryonic forebrain beginning at E11.5(Goebbels et al., 2006). These animals survive to adulthood, allowing thorough assessment of their behaviour. A battery of tests revealed that *Atrx*^NEXCre^ male and female mice have impaired fear memory as well as ASD features. We also observed sexually dimorphic behaviour differences, including male-specific aggressivity and altered sensory gating and female-specific social avoidance. Overall, these mice represent a clinically relevant model to study the underlying causes of ASD and ID caused by ATRX deficiency in neurons.

## RESULTS

### Generating mice with ATRX deletion in excitatory neurons of the embryonic forebrain

Mice were generated that lack *Atrx* in forebrain post-mitotic excitatory neurons starting at embryonic day (E)11.5, using the NEX-Cre driver line of mice (Goebbels et al., 2006). Neuronal helix-loop-helix protein-1 (NEX, also called NeuroD6) is a transcription factor expressed in post-mitotic immature neurons of the dorsal forebrain and is involved in terminal differentiation (Ross et al., 2003). We confirmed loss of ATRX by immunohistochemistry of coronal brain cryosections at E13.5 and P20. The results show that by E13.5, loss of ATRX is apparent in the cortical plate where neurons begin differentiation and express βIII-tubulin (**Fig. 1A,B**). At P20, ATRX staining demonstrates that the protein is absent in NeuN-positive neurons of *Atrx*^NEXCre^ male cortex and hippocampus (**Fig. 1C,D**). We note that ATRX protein is still expressed in a subset of cells in the dentate gyrus of the hippocampus at P20 and 3 months, likely corresponding to neuroprogenitor cells and GABAergic interneurons (Dieni et al., 2013). Confirmation of ATRX deficiency was also confirmed in *Atrx*^NEXCre^ female brain cryosections at E13.5 and P20 (**Fig. S1 A-D**).

**Figure 1:**
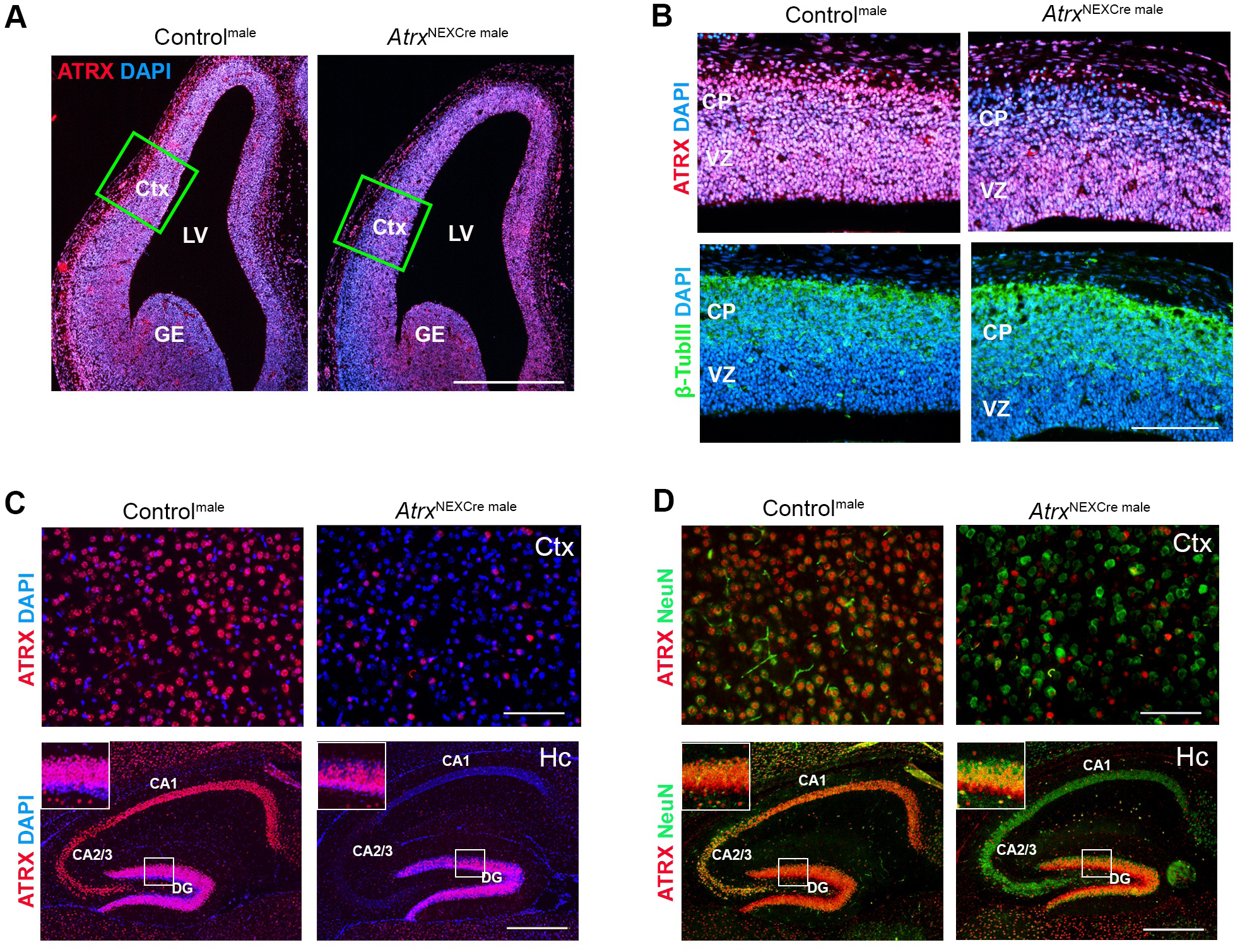
Validation of *Atrx* deletion in *Atrx*^NEXCre^ male mice. **A)** ATRX immunofluorescence staining of E13.5 control and *Atrx*^NEXCre^ brain coronal cryosections in red. DAPI counterstaining is shown in blue. Scale bar, 570 µm. **B)** Magnified view of ATRX (top, red) and βIII-tubulin (bottom, green) staining in the developing cortical plate. Scale bar, 144 µm. **C)** ATRX (red) and DAPI (blue) staining in the cortex and hippocampus of *Atrx*^NEXCre^ and control mice at P20. **D)** ATRX (red) and NeuN (green) staining in the cortex and hippocampus at P20. Cortex scale bar,100 µm and hippocampus scale bar, 400 µm. CA: cornu Ammonis, CP: cortical plate, Ctx: cortex, DG: dentate gyrus, GE: ganglionic eminence, LV: lateral ventricle, VZ: ventricular zone.

### Learning and memory impairments in *Atrx*^NEXCre^ mice

Learning and memory paradigms are used in animal models to measure cognitive deficits that compare to intellectual disabilities clinically. Two common learning and memory paradigms used to evaluate mouse models of ID are the contextual fear memory test and the Morris water maze paradigm (Mamiya et al., 2009; Vorhees & Williams, 2006). First, we utilized the contextual fear memory test, where mice are placed in an opaque box with distinct visual patterns on the side and a metal grate flooring for 3 minutes during the conditioning portion of the test. At the 2.5 minute mark, animals receive a foot-shock through the metal grate floor. In response to the shock, they exhibit freezing behaviour indicating fear (**Fig. 2A**). In the memory portion of the test, animals are placed back in the same apparatus either 1.5 or 24 hours post foot-shock, and freezing behaviour is measured as indication of fear memory. We found that male and female *Atrx*^NEXCre^ mice spend significantly less time freezing compared to control mice at 1.5 hours (Male: p<0.0001, F=1.299; Female: p<0.0001, F=3.250) and 24 hours post foot shock (Male: p=0.0002, F=2.019; Female: p<0.0001, F=2.830) (**Fig. 2B,C**), indicating short- and long-term memory deficits. Discrimination indices revealed no sex-difference in fear memory due to loss of ATRX (**Fig. 2B,C**).

**Figure 2:**
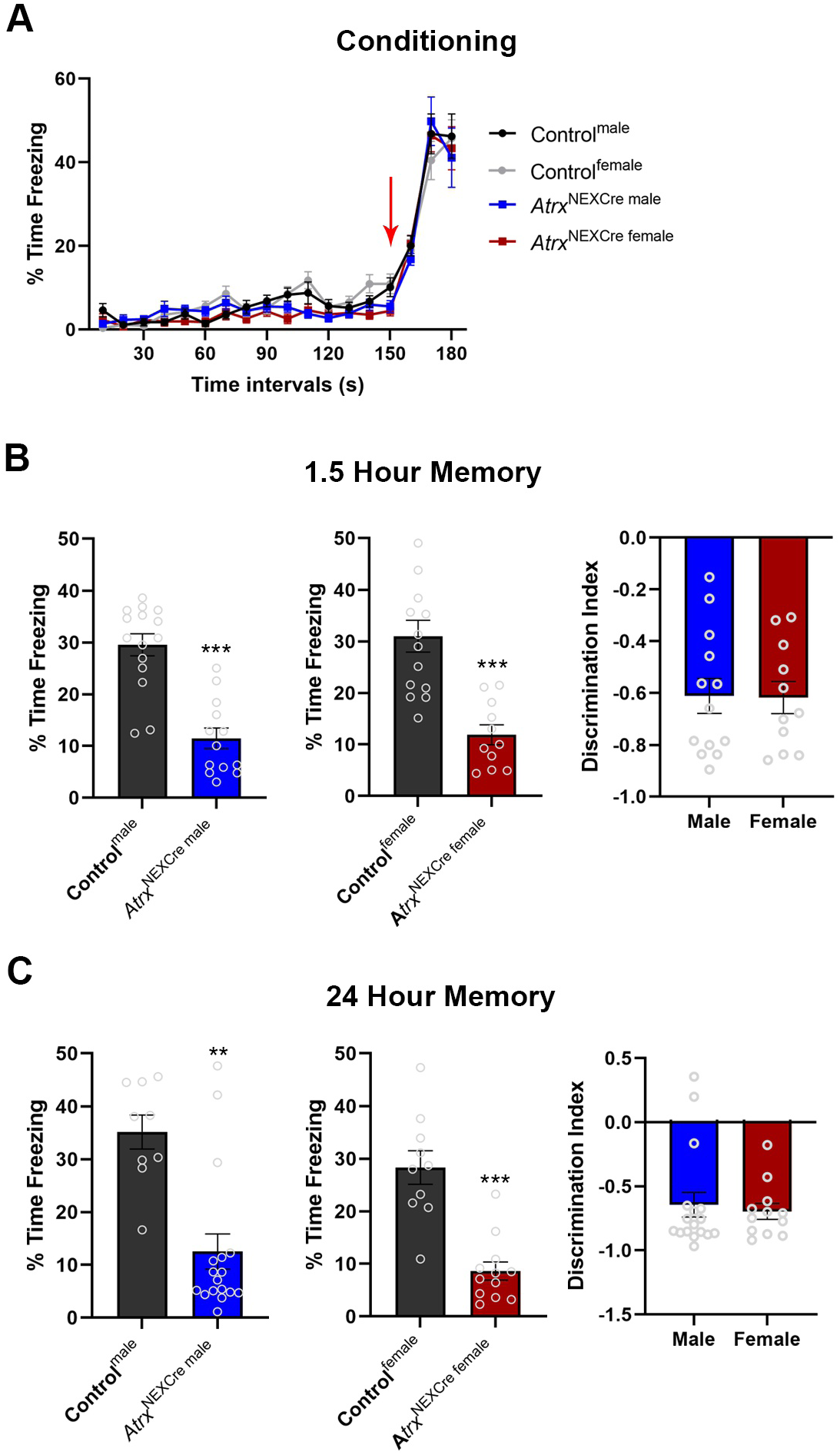
Contextual fear memory deficits in male and female *Atrx*^NexCre^ mice. **A)** 3-month-old mice were placed in contextual fear apparatus for 3-minutes and received a foot-shock at 2.5-minutes (red arrow). Mice of each sex and genotype as indicated responded with increased freezing after the foot-shock. **(B,C)** *Atrx*^NEXCre^ mice spent significantly less time freezing compared to control mice at 1.5 and 24 hours after foot shock (1.5 hours male: p<0.0001, F=1.299, Control^male^ n=15, *Atrx*^NEXCre^ male n=13. 1.5 hours female: p<0.0001, F=3.240, Control^female^ n=14, *Atrx*^NEXCre^ ^female^ n=11. 24 hours male: p=0.0002, F=2.019, Control^male^ n=9, *Atrx*^NEXCre^ ^male^ n=17. 24 hours female: p<0.0001, F=2.830, Control^female^ n=10, *Atrx*^NEXCre^ ^female^ n=12). Discrimination indices were calculated for male and female mice to determine the effect of loss of ATRX on freezing behaviour and showed no sex difference in fear memory deficits. Error bars represent +/-SEM (Students t-test, ** p<0.001, ***p<0.0001).

For the Morris water maze, mice are placed into a water bath and trained over 4-days to find a submerged platform using spatial cues (Vorhees and Williams, 2006). In this test, male and female *Atrx*^NEXCre^ mice failed to find the platform, even after 4-days of training (**Fig. S2A**). As the training days continued, the *Atrx*^NEXCre^ mice spent less time swimming and tended to remain close to the wall or displayed floating behaviour (**Fig. S2B, Movie 1**). To explore whether these behaviours reflect increased apathy, we performed a 6-minute forced swim test, where animals are placed in a tall cylinder of water for 6-minutes and time immobile is measured (Can et al., 2011). However, *Atrx*^NEXCre^ mice did not exhibit decreased time swimming compared to sex-matched control mice, ruling out this possibility (**Fig. S2C**). While the data on memory deficits remains inconclusive, it is noteworthy that the *Atrx*^NEXCre^ mice display atypical swimming behaviour and a lack of participation in the test, which is consistent across both sexes.

### Decreased anxiety and autism-associated behaviours in *Atrx*^NEXCre^ mice

Hyperactivity disorders are often co-diagnosed in ASD patients and ASD mouse models (Luo et al., 2018; Simonoff et al., 2008). We therefore tested the activity levels and observed that male and female *Atrx*^NEXCre^ mice display hyperactivity as measured by distance travelled and velocity in an open field (Male: Distance p=<0.0001, F=6.974, Velocity p=0.0009, F=3.678; Female: Distance p=<0.0001, F=13.02, Velocity p=0.0012, F=4.256)(**Fig. 3A,B**).

**Figure 3:**
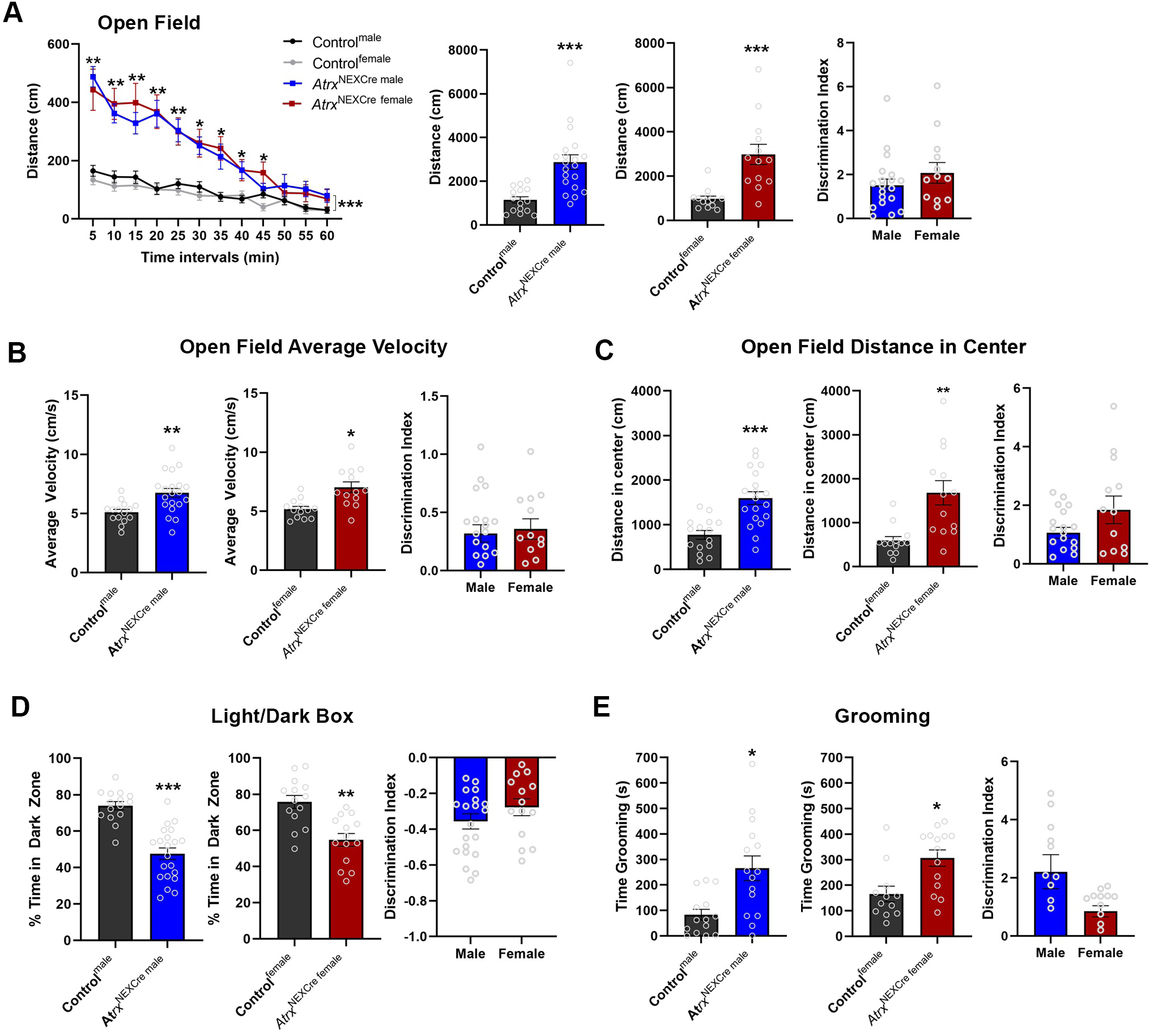
*Atrx*^NEXCre^ mice are hyperactive, less anxious and overgroom compared to control mice. **A)** Open field testing of *Atrx*^NEXCre^ male and female mice shows significantly increased locomotor movement compared to controls as seen over-time in 5-minute intervals and total distance travelled in 1 hour (Male: p<0.0001, F=6.974, Control^male^ n=15, *Atrx*^NEXCre^ ^male^ n=20; Female: p<0.0001, F=13.02, Control^female^ n=13, *Atrx*^NEXCre^ ^female^ n=13). **B)** Loss of ATRX in excitatory neurons results in significantly increased velocity of movement (Male p=0.0009, F=3.678, Female p=0.0012, F=4.256) and **C)** decreased distance travelled in the center of the open-field (Male p<0.0001, F=2.691, Female p=0.0004, F=9.146). **D)** Time spent in the dark zone in the light-dark box paradigm (Male: p<0.0001, F=2.746, Control^male^ n=15, *Atrx*^NEXCre^ ^male^ n=21, Female: p=0.0003, F=1.016, Control^female^ n=14, *Atrx*^NEXCre^ ^female^ n=14). **E)** Adult *Atrx*^NEXCre^ mice spend more time grooming than control mice (Male: p=0.0030, F=5.450, Control^male^ n=18, *Atrx*^NEXCre^ ^male^ n=15, Female: p=0.0042, F=1.016, Control^female^ n=12, *Atrx*^NEXCre^ ^female^ n=14). Error bars represent +/-SEM. Students t-test, Mann Whitney, or Two-way Anova with Sidak post-hoc (* p<0.05, ** p<0.001, ***p<0.0001).

Travel in the center of the open field is used as a measure of anxiety levels. *Atrx*^NEXCre^ male and female mice travelled significantly more distance in the center of the open field compared to sex-matched controls (Male: p<0.0001, F=2.691, Female: p=0.0012, F=4.256) suggesting that loss of neuronal ATRX has anxiolytic effects (**Fig. 3C)**. To further support these findings, *Atrx*^NEXCre^ mice were evaluated in the light-dark box paradigm for 15-minutes, and the amount of time spent in the open and light area compared to the enclosed dark space was measured. Both male and female *Atrx*^NEXCre^ mice spent significantly less time in the dark area compared to controls (Male: p<0.0001, F=2.746, Female: p=0.0003, F=1.016), confirming lowered anxiety in these mice (**Fig. 3D)**. Discrimination indices show no sex differences in this phenotype (**Fig. 3A-D**). We also attempted to evaluate anxiety utilizing the elevated plus maze, however the *Atrx*^NEXCre^ mice jumped off the apparatus and could not complete the task.

Stereotypies represent a core autistic feature (Abrahams and Geschwind, 2008) that is apparent by an overgrooming phenotype in mice. Grooming behaviour was evaluated in control and *Atrx^NEXCre^* mice over a 15-minute period. This revealed that male and female *Atrx^NEXCre^* mice display repetitive behaviours, as shown by increased time spent grooming (Male: p=0.0030, F=5.450, Female: p=0.0042, F=1.281) (**Fig. 3E**). Repetitive grooming in some mice led to large patches of fur missing, and in severe cases mice needed to be euthanized for self-injuries (**Fig. S3A**). Statistical testing of the discrimination ratio for increased grooming behaviour indicates no significant changes between sexes.

### Male-specific sensory gating deficit in *Atrx*^NexCre^ mice

Sensory processing abnormalities are implicated in patients with ASD. To test this phenotype, we utilized a pre-pulse inhibition test to examine startle response and sensory gating (Davis, 1984; Martin-Kenny and Bérubé, 2020). First, animals were exposed to 50 trials of a 115 db pulse, lasting 20 ms with 20 s between each trial. We observed an exaggerated startle response to this acoustic stimulus in male, but not female *Atrx*^NexCre^ mice (Male: p=0.0008, F=18.08)(**Fig. 4A**). Next, animals were exposed to the pre-pulse inhibition portion of the test, consisting of a pulse only at 115 db, 40 ms in length, and four types of pre-pulse trials: Type 1: 75 db, 30 ms prior to the startle pulse, Type 2: 80 db, 30 ms prior to startle, Type 3: 75 db, 100 ms prior to startle, and Type 4: 80 db, 100 ms prior to startle. Male *Atrx*^NEXCre^ mice displayed decreased pre-pulse inhibition compared to controls (Male: Type 1: p=0.0286, F=1.414, Type 2: p=0.0261, F=2.048, Type 3: p=0.0042, F=1.549, Type 4:p=0.0008, F=2.510). Conversely, female *Atrx*^NEXCre^ mice did not show a similar reduction in PPI (**Fig. 4B,C**). Statistical analysis reveals a significant sex difference in Types 1, 2 and 4 PPI settings [Type 1 (p=0.0125, F=2.955), Type 2 (p=0.0074, F=6.064), and Type 4 (p=0.0093, F=1.338)]. Together these data reveal a male-specific effect of ATRX loss on sensory gating.

**Figure 4:**
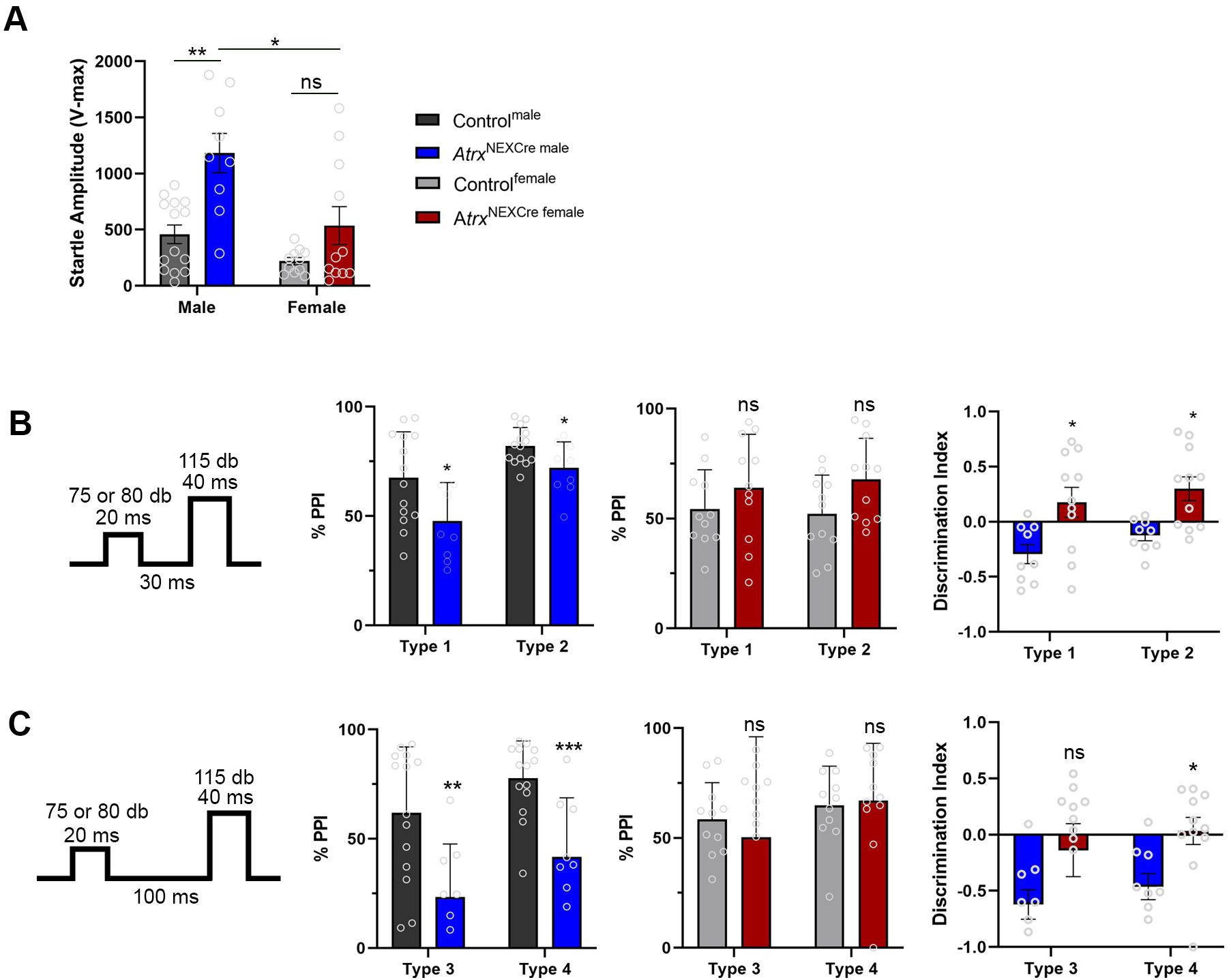
*Atrx*^NexCre^ mice exhibit sex-specific deficits in sensory gating. **A)** *Atrx*^NEXCre^ male mice display an exaggerated auditory startle response compared to controls (p=0.0008, F=18.08), whereas *Atrx*^NEXCre^ females do not. **B)** *Atrx*^NEXCre^ male mice exhibit decreased pre-pulse inhibition for Type 1 (30ms, 75db) (p=0.0286, F=1.414), Type 2 (30ms, 80db) (p=0.0261, F=2.048), **C)** Type 3 (75db, 100ms) (p=0.0042, F=1.549) and Type 4 settings (80db, 100ms) (p=0.0008, F=2.510). Control^male^ n=14, *Atrx*^NEXCre^ ^male^ n=9, Control^female^ n=11, *Atrx*^NEXCre^ ^female^ n=11. Sex differences were detected in Type 1 (p=0.0125, F=2.955), Type 2 (p=0.0074, F=6.064), and Type 4 settings (p=0.0093, F=1.338). Error bars represent +/-SEM. Students t-test, Mann Whitney, or Two-way Anova with Sidak post-hoc performed when appropriate (* p<0.05, ** p<0.001, ***p<0.0001).

### Social behaviour deficits in *Atrx*^NEXCre^ mice

The three-chamber test was used to evaluate social phenotypes. All experimental animals freely explored the chamber during a 10-minute habituation period (**Fig. S3B**). *Atrx*^NEXCre^ female mice failed to show the expected preference for the stranger mouse over the object in this test (**Fig. 5A**), while the *Atrx*^NEXCre^ males behaved similarly to control mice. Based on the sociability index, female (but not male) *Atrx*^NEXCre^ mice have decreased sociability (Female: p=0.0120, F=2.527) (**Fig. 5B**). However, discrimination index analysis revealed no significant sex difference in sociability in *Atrx*^NEXCre^ mice. We noticed that *Atrx*^NEXCre^ males were exhibiting a tail rattling phenotype when in the presence of the stranger mice during the three-chamber test. Tail rattling is typically identified as an aggressive behaviour in mice, often associated with fear or to show threat in aggressive social encounters(Scott, 1966). Quantification shows that *Atrx*^NEXCre^ male mice exhibit significantly more tail rattling events at the stranger mouse compared to controls (p=0.0013, F=10.35) and that female mice do not exhibit this behaviour (**Fig. 5C**). These results identify a male-specific increase in social aggression in *Atrx*^NEXCre^ mice.

**Figure 5:**
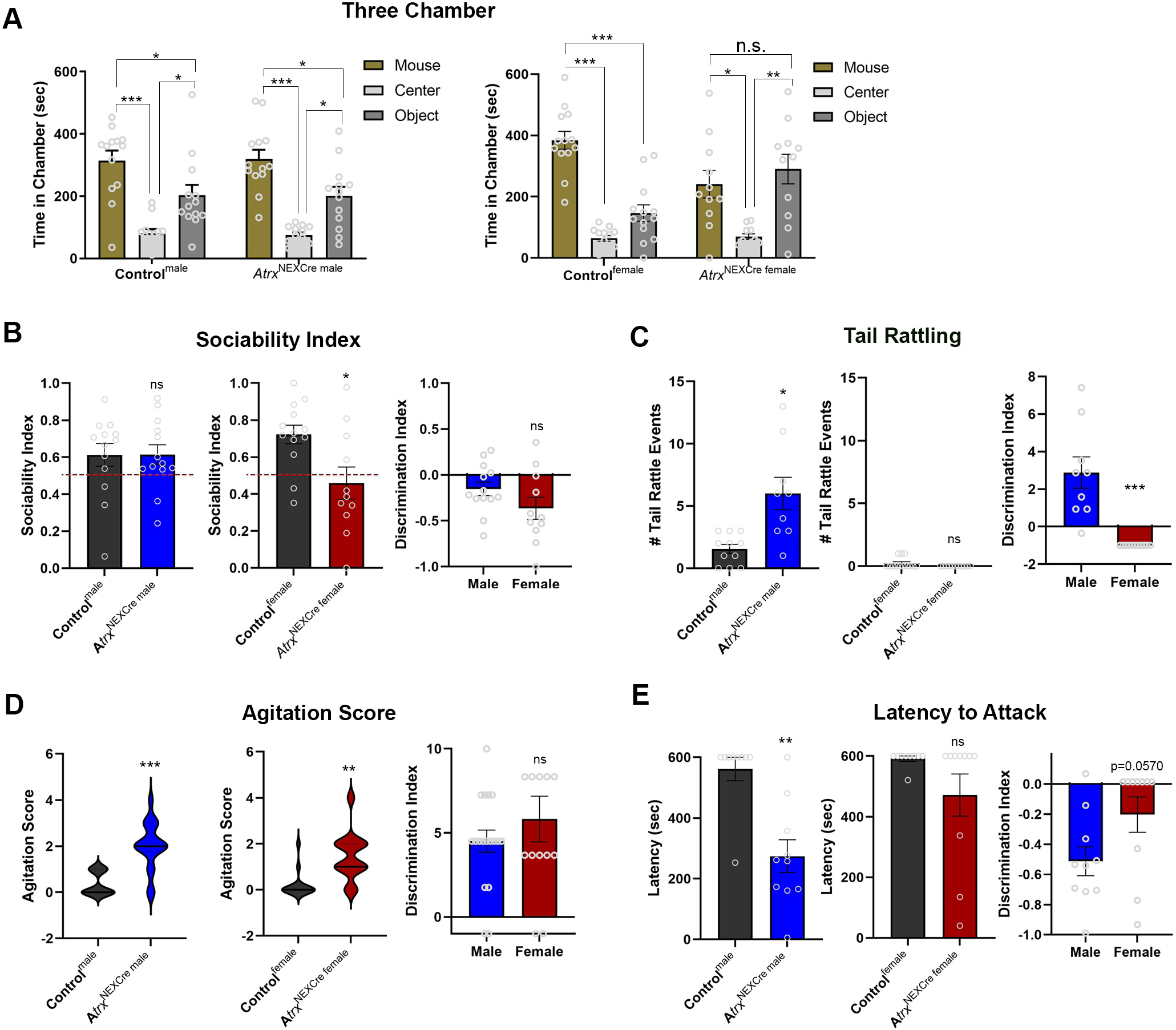
*Atrx*^NEXCre^ mice exhibit changes in social behaviour. **A)** Time spent by mice of the indicated sex and genotype in each chamber of the three chamber test (Control^male^ p=0.0098, *Atrx*^NEXCre^ ^male^ p=0.0058, Control^female^ p<0.0001). **B)** Male *Atrx*^NEXCre^ mice show no difference in sociability compared to controls in the three-chamber test (Control^male^: n=13, *Atrx*^NEXCre^ ^male^ n=13), whereas female *Atrx*^NEXCre^ mice display decreased sociability (Female: p=0.0120, F=2,527, Control^female^ n=13, *Atrx*^NEXCre^ ^female^ n=11). **C)** *Atrx*^NEXCre^ male mice exhibit significantly more tail rattling events in the presence of a stranger mouse (p=0.0013, F=10.35), while *Atrx*^NEXCre^ female mice do not exhibit this behaviour. **D)** *Atrx*^NEXCre^ male and female mice are more agitated upon handling, as determined by an elevated agitation score (Male: p<0.0001, F=4.160, Control^male^ n=11, *Atrx*^NEXCre^ ^male^ n=18, Female: p=0.0003, F=3.290, Control^female^ n=14, *Atrx*^NEXCre^ ^female^ n=13). **E)** *Atrx*^NEXCre^ male mice exhibit decreased latency to attack in the juvenile resident intruder paradigm, and *Atrx*^NEXCre^ female mice do not (Male: p=0.0006, F=2.187, Control^male^ n=9, *Atrx*^NEXCre^ ^male^=10, Female: Control^female^ n=9, *Atrx*^NEXCre^ ^female^ n=10). Error bars represent +/-SEM. Two-way Anova with Sidak post-hoc, Students t-test, or Mann Whitney performed when appropriate (*p<0.05, ** p<0.001, ***p<0.0001).

Impaired social memory is commonly observed in patients with ASD. In mouse models, social memory can be assessed during the third part of the three-chamber test, which involves a social novelty test. However, we were unable to use social memory data from our three-chamber tests, as loss of ATRX significantly affects the olfaction of both male and female *Atrx*^NEXCre^ mice, and mice rely on olfaction for social novelty (**Fig. S3B**).

We observed that *Atrx*^NEXCre^ mice are agitated in the presence of a handler. We quantified this unusual behaviour using an agitation score (0-5) that reflects biting, trunk curl and vocalization events during handling (**Movie 2**). According to these criteria, both *Atrx*^NEXCre^ male and female mice are significantly more agitated upon handling compared to controls (Male: p<0.0001, F=4.160, Female: p=0.0003, F=3.290) (**Fig. 5D**). No sex differences were identified for the agitation score. We also found that *Atrx*^NEXCre^ male mice were aggressive towards their littermates, occasionally resulting in separation due to injuries (8/35 cages). Issues with littermates fighting was not noted with *Atrx*^NEXCre^ female mice. To establish the extent of this aggressive behaviour, we performed a juvenile resident intruder assay. The results show that *Atrx*^NEXCre^ male mice have a decreased latency of attack, showing a significant increase in aggression not seen with *Atrx*^NEXCre^ female mice (Male: p=0.0006, F=2.187) (**Fig. 5E**). The discrimination index indicates no significant sex difference in their response to the intruder (**Fig. 5E**). These data suggest that loss of ATRX embryonically in excitatory neurons affects social communication in adult male and female mice, with a more aggressive social behaviour in males, and a decreased sociability in females.

## DISCUSSION

The main goal of this study was to determine if embryonic deletion of ATRX in forebrain excitatory neurons would result in a broader spectrum of deficits than previously observed when ATRX is deleted postnatally in these same neurons. When ATRX is deleted postnatally in excitatory neurons, the mice display impaired long term spatial memory without ASD-like features (Martin-Kenny and Bérubé, 2020; Tamming et al., 2020). However, a subset of ATR-X syndrome patients exhibit more widespread cognitive and neurological problems (Gibbons and Higgs, 2000). Therefore, we hypothesized that earlier deletion of *Atrx* was required to induce ASD-associated traits. Indeed, we observed the two core ASD features in adult *Atrx*^NEXCre^ mice: social deficits and repetitive behaviours. We also observed hyperactivity and a male-specific deficit in acoustic sensory gating. According to the DSM-V (American Psychiatric Association, 2013), impaired sensory processing is now considered a core feature of ASD. Based on the published and current models of ATRX inactivation in neurons, it appears that there is a period of vulnerability (mid-gestation to early adolescence) where the dorsal forebrain is highly sensitive to ATRX loss of function, with dire consequences on behaviour in adulthood, particularly for autistic traits. This is supported by previous reports showing that ASD risk genes are expressed at high levels in the cortex in mid-to-late fetal development and largely enriched in maturing excitatory neuron lineages (Satterstrom et al., 2020). In addition, studies show that patients with ASD have atypical brain development, with changes in proper neuronal migration in the cortex (Hutsler et al., 2007) and many transcription factors that appear in mid-gestation and play an important role in corticogenesis have been shown to be ASD-risk genes (Fazel Darbandi et al., 2020; Kwan, 2013).

The *Atrx*^NEXCre^ mice characterized in this study offer a good model to help understand the structural and molecular changes that occur early during forebrain development and that lead to altered cognitive abilities. Research focusing on mouse models with mutations in synaptic proteins have found similar behavioural alterations to *Atrx*^NEXCre^. For examples, mice with mutations in *Shank3*, a gene encoding a postsynaptic protein, exhibit hyper-reactive behaviours, increase in escape tendencies, aggression, and repetitive behaviours (Drapeau et al., 2018; Zhang et al., 2021). Animal models containing mutations in the autistic linked post-synaptic neuroligin family of proteins display hyperactivity, repetitive behaviours and aggression (El-Kordi et al., 2013; Kohl et al., 2015; Rothwell et al., 2014). Previously, our lab identified several ATRX target genes related to synaptic function, including *Neuroligin 4* (*Nlgn4*) that functions at the post-synaptic cleft and is a known autism-associated gene (Jamain et al., 2003; Levy et al., 2015). Other relevant ATRX-regulated genes include miRNA137, whose target genes are involved in proper synapse function (Tamming et al., 2020) and *Xlr3b*, an imprinted gene involved in the regulation of mRNA transport in dendrites (Shioda et al., 2018). Future work should focus on characterizing the transcriptional changes in the male and female *Atrx*^NEXCre^ mice during the identified period of vulnerability to gain mechanistic understanding of the ensuing cognitive dysfunction.

We observed significant short- and long-term deficits in contextual fear memory in *Atrx*^NEXCre^ mice of both sexes. This contrasts with *Atrx*^CamKIICre^ mice (deletion starting at ∼P20), in which fear memory deficits were only observed in males (Tamming et al., 2020). Female protection theories have been extensively reviewed and suggest that estrogen levels play a neuroprotective role in preventing cognitive impairments (Azcoitia et al., 2019; Enriquez et al., 2021). A previous study used a computational model to investigate molecular networks of copy number variants (CNVs) in both male and female patients with ASD. They found that CNVs in females affect a larger number of genes, and that genes disrupted in female ASD patients were functionally more important in molecular networks. These data support previous hypotheses suggesting that females need to experience a harsher effect on connectivity than males to experience ASD symptoms (Gilman et al., 2011). The difference in this protective effect between the *Atrx*^CamKIICre^ and *Atrx*^NEXCre^ female mice suggests that in a neuroplastic setting, hormones can impart neuroprotective effect; however, in the case of an embryonic deletion, the anomalous network connectivity might not be as malleable.. Investigation into the sex-specific cellular and connectivity changes in *Atrx*^NEXCre^ mice could shed light on protective mechanisms or reversibility of the observed behaviours.

In our model, both male and female *Atrx*^NEXCre^ mice exhibit hyperactivity and repetitive behaviours. However, social communication alterations differed, with male *Atrx*^NEXCre^ mice experiencing social aggression and female *Atrx*^NEXCre^ mice social avoidance. Finally, we observed that acoustic hypersensitivity and decreased PPI were male specific. Clinically, males are biased in diagnosis of many NDDs, including ID, ASD, and ADHD. The reason, whether female protection, male vulnerability, diagnosis bias, or societal differences between males and females is highly debated (Ferri et al., 2018; Rutherford et al., 2016). Within the scientific literature, there is an increasing concern regarding the under identification of ASD in females when compared to males (Kilmer & Boykin, 2022). Presently, there is a dearth of comprehensive research that encompasses both male and female mice in their investigations. Consequently, it is unclear how females are affected compared to males, impeding our understanding of the underlying reasons for underdiagnosis. Acquisition of sex-specific data in ASD could prove instrumental in elucidating this phenomenon.

### Final Conclusions

The behavioural characterization of the *Atrx*^NEXCre^ mouse model establishes it as a viable system for studying a range of neurological abnormalities associated with ATRX dysfunction. This model holds potential in unraveling the mechanisms related to the vulnerable period for ASD during brain development, as well as shedding light on sex differences in ASD and theories of female protection. More work is required to identify potential connectivity disruptions, downstream targets affected by ATRX loss, and the overall molecular and structural distinctions between male and female *Atrx*^NEXCre^ mice. Ultimately, this model provides a clinically relevant platform to identify therapeutic targets and serves as a preclinical model for testing potential therapies.

## MATERIALS AND METHODS

### Animal care and husbandry

All experiments were performed using *Mus musculus*, generated with conditional inactivation of *Atrx* in post-mitotic excitatory neurons starting at E11.5, by crossing 129SV female mice heterozygous for *Atrx*^loxP^ sites (Berube et al., 2005) with C57BL/6 male mice expressing Cre recombinase under the control of NEX-Cre gene promotor (Neurod6^tm1(cre)Kan^) (Goebbels et al., 2006). Male progeny with a floxed *Atrx* allele on the X chromosome and the transgenic NEXCre allele are ATRX-null (*Atrx*^NEXCre^ ^male^). Control male and female mice only have the NEXCre allele (Control^male^, Control^female^). To obtain homozygous *Atrx*^loxP^ females, 129SV female mice heterozygous for *Atrx*^loxP^ sites are crossed with C57BL/6;129SV male mice expressing NEX-Cre and have a floxed *Atrx* allele, offspring homozygous for *Atrx*^loxP^ expressing NEXCre are knockouts (*Atrx*^NEXCre^ ^female^). Mice were exposed to a 12-hour-light/12-hour-dark cycle with water and chow ad libitum. All procedures involving animals were conducted in accordance with the regulations of the Animals for Research Act of the province of Ontario and approved by the University of Western Ontario Animal Care and Use Committee (2021-049).Ear notch biopsies were taken at P8-10 for animal ID and genotyping purposes. Genotyping was performed as previously described (Berube et al., 2005) using the primers in **Table 1**.

**Table 1:**
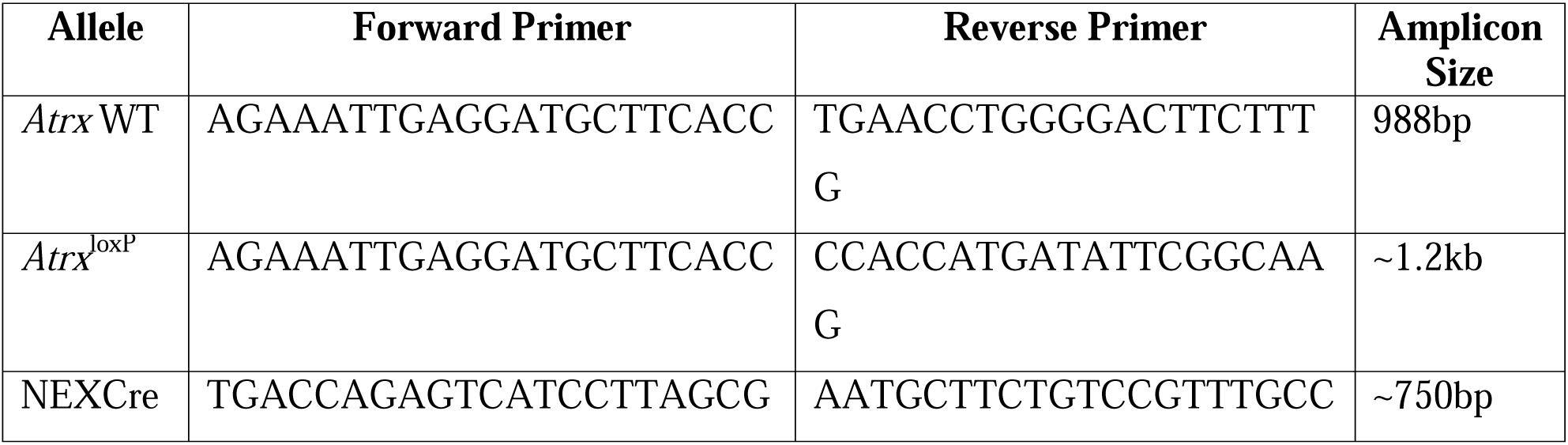
Genotyping Primers.

### Immunofluorescence and microscopy

For histological analysis of embryonic samples, timed matings were arranged late afternoon and the following midday, day of vaginal plug discovery, was considered embryonic day (E)0.5. At E13.5 pregnant females were anesthetized by CO2, sacrificed by cervical dislocation, and embryos were dissected and placed into 4% PFA overnight for fixation. For histological analysis at P20, mice were perfused with cold 1xPBS, followed by perfusion with cold 4% PFA overnight. After fixation all samples were washed with PBS and put into 30% sucrose for a minimum of one week prior to embedding in Cryomatrix™ (Epredia cat#6769006) in an ‘ethanol-dry ice slurry’, and then placed at −80°C for storage. Samples were sectioned at 10µM thickness using a Leica cryostat (CM3050 S). Immunostaining was performed by rehydration in 1xPBS followed by an antigen retrieval by boiling in sodium citrate pH 6 containing 0.5% tween for 15 minutes. A permeabilization step consisting of a 10 min wash with 1xPBS containing 0.03% Triton X was followed by 1-hour of blocking in 1xPBS, 0.3% Triton X, 5% Goat serum. Primary antibodies were added in blocking solution [ATRX (Santa Cruz sc15408): 1:150 (Rb), βIII-tubulin (Biolegend 801201) 1:2000 (Ms), NeuN (Millipore MAB377): 1:150 (ms), PV (Sigma P3088): 1:1000 (Ms)] overnight at 4°C, followed by washing with 1xPBS, 0.3% Triton X solution. Secondary antibodies were added as appropriate (Thermo Scientific 1:1000 Alexa Fluor 488 anti-mouse cat#A11017, Alexa Fluor 594 anti-mouse cat#A11032, Alexa Fluor 594 anti-rabbit cat#A11012, Alexa Fluor 488 anti-rabbit cat#A11034) for 1 hour, slides washed with 1XPBS 0.3% Triton X, treated with a DAPI stain, washed, and mounted using ImmunoMount (Thermo Scientific cat# 9990402). Stained cryosections were imaged on a Leica microscope (CTR 6500) using Open-lab and Velocity version 6.0.1 software for analysis.

### Behaviour Testing

Behavioral assessments were performed at 3-6 months of age, starting with less demanding tasks (open field tests, light-dark paradigm) to more demanding ones (contextual fear, resident intruder). ARRIVE guidelines were followed: animal groups were randomized, experimenters were blind to the genotypes, software-based analysis was used to score performance in all possible tasks. When software-based analysis was not possible, video recordings were coded and revealed to the experimenter after analysis. We have found that sample sizes of 10-15 per genotype is large enough to reach statistical significance in behavioural paradigms (Martin-Kenny and Bérubé, 2020; Tamming et al., 2017; Tamming et al., 2020). All behavioral experiments were performed between 9:00 AM and 4:00 PM.

#### Open-Field

Mice were placed in a 20 cm x 20 cm open arena with 30 cm high walls and left to explore for 1 hour. Animals were brought into the testing room 30 minutes prior to the start of the test to habituate to the room in their home cages. Locomotor activity, velocity, and distance travelled in the center of the open-field were all recorded in 5-minute intervals by AccuScan Instrument (Cogram et al., 2019; Tamming et al., 2017).

#### Light/Dark Box

Mice were placed in a 40 cm x 40 cm arena with 30 cm high walls, and half the arena had a dark-box insert with a small open-slot where they could freely move back and forth between the light and dark areas. Animals were brought into the testing room in their home cages 15 minutes prior to the start of the test for habituation. Animals were placed in the light area of the apparatus and left to freely explore for 15-minutes. Time spent in the light-zone and dark-zone was measured in 5-minute intervals by AccuScan Instrument (Cogram et al., 2019).

#### Grooming

Animals were individually habituated in an empty cage (home cage with no bedding) for 15 minutes prior to the start of the test. Mice were then recorded for 15 minutes using the Anymaze video-tracking system. The time spent grooming was manually scored from the videos(Stoodley et al., 2017).

#### Olfaction

The odor habituation and discrimination assay was performed as previously described (Arbuckle et al., 2015). Briefly, animals were habituated in a clean cage with a wire lid for 30 minutes prior to the start of the test. Mice were then presented with a cotton swab with either water, almond or banana flavouring (1:100 dilution of Club House extract). Each scent was presented for three sequential 2-minute trials and the time spent sniffing the odour was recorded. Sniffing was defined as the animal’s nose oriented towards and near the cotton swab (2 cm or closer).

#### Three-Chamber

The three-chamber apparatus was used to observe sociability, as previously described (El-Kordi et al., 2013) with minor modifications. Animals were habituated to the room for 20 minutes prior to beginning the assessment, and then allowed a 10-minute habituation trial where they freely explored the three-chamber apparatus containing wire cages to be used in the test trial. For the sociability test trial, on one side of the three-chamber a same-sex and age-matched wild-type C57Bl/6 stranger mouse was placed under the wire cage, and on the opposite side an object was placed under an identical wire cage. The test animal was then allowed to once again freely explore the apparatus and time spent in each chamber was recorded by AnyMaze video-tracking system. The sociability index was calculated as follows time with stranger /(time with stranger + time with object) x 100. Tail rattling events were obtained from the same videos (Cogram et al., 2019).

#### PPI

To assess sensory gating, we performed a startle response and pre-pulse inhibition test, as previously described (Davis, 1984; Martin-Kenny and Bérubé, 2020) (SR-LAB, San Diego Instruments). Briefly, animals were placed in the chamber apparatus and exposed to background noise (65 db) for 5-minutes for two habituation days prior to the test day. On test day, animals were again placed in the chamber and underwent a 10-minute acclimation with background noise (65 db), followed by a habituation block, which consisted of fifty acoustic startle pulses (115 db, 20 ms in length) at 20 second intervals. Finally, animals were exposed to a pre-pulse inhibition block which consisted of ten sets of five trial types randomly ordered with variable intervals of 10, 15 or 20 seconds between each trial. Four of the five trial types had a pre-pulse (75 or 80 db, 20 ms in length) 30 ms or 100 ms prior to the startle stimulus (115 db, 40 ms in length). The fifth trial type was the acoustic startle stimulus alone (115 db, 40 ms in length). The startle response represents the movement the platform, generating a measurable force by the SR-LAB software. The average startle response of the ten trials for trial type was calculated, and pre-pulse startle responses were normalized to the startle alone trial.

#### Contextual Fear

Contextual fear memory was measured utilizing a 20 cm x 10 cm acrylic enclosure, as previously described (Tamming et al., 2020) with minor modifications. The enclosure was wrapped in white paper, with a distinct striped wall on one side, a large black star on the opposite wall, and a metal grid floor. The metal grid floor was equipped with an electric shock generator, and a video recording was taken from above the apparatus and recorded using AnyMaze video tracking software. During the conditioning portion of the test, animals could freely explore the enclosure for 3-minutes. At the 150 second mark the animals were given a shock (2 mA, 180 V, ∼1 second in length) through the metal grated floor and left to explore the cage another 30 seconds before returning to their home cage. Either 1.5 or 24-hours after receiving the shock, animals were placed back into the enclosure for 6-minutes and time spent freezing was measured. Time spent freezing is defined as complete immobility.

#### Morris Water Maze

The Morris water maze task was performed to determine alterations in spatial learning in memory, as previously described (Tamming et al., 2020; Vorhees and Williams, 2006) with minor modifications. The maze is a 1.5 m diameter pool with a water temperature of 26°C, and black and white shapes act as spatial cues on each of the four walls surrounding the pool, creating four quadrants. Animals were given four trials (90 s) a day for four consecutive days, with a 15-minute intertrial period, to find a submerged platform (1.5 cm below the water surface). If the animals did not find the platform within the 90-second trial, they were gently guided to the platform. On the fifth day, the platform was removed, and animals were given one 60-second test trial, with time spent in each quadrant recorded. All training trials and test-day trials were recorded, and movement monitored using AnyMaze tracking software.

#### Forced Swim Test

To test if mice displayed apathy in water, the forced swim test was performed (Can et al., 2011). Briefly, animals were placed in a beaker (15 cm diameter) of water (26°C, depth of 9 cm) for six-minutes. The first two minutes of the test was a habituation period, a camera positioned above the beaker recorded the last four-minutes of the test in AnyMaze software. Time spent immobile was determined during the four-minute test period. Time immobile refers to time floating with no tail or paw movement in the water. Small movements required to keep afloat was considered as time immobile.

#### Agitation score

We developed an agitation score based on handling during low-stress conditions. Three separate factors were combined to make a score from 0-5. First, a biting score was given (0-3) based on the number of times the handler was bitten (while wearing a protective glove) on three different days. Next, a 15-second video was taken moving the animal from one housing cage to another, and during the transfer the animal has held by the base of the tail for ∼8 seconds. A trunk curl score (0=absent, 1=present) during 8 seconds of being held by the tail, and vocalization (0=absent, 1=present) during the 15 second video were also added for the total agitation score.

#### Resident Intruder

To assess aggression a resident-intruder paradigm was used, as previously described (El-Kordi et al., 2013)with minor alterations. Experimental animals were separated and housed individually for 21 days prior to test day. Juvenile (8-10weeks of age) same-sex group-housed mice were used as intruders and were ∼30-35% smaller in weight than experimental animals. Each intruder was only used once to avoid winner or loser effects. Intruders were introduced into the resident cage (at least a couple days dirty) and video tracking using AnyMaze software was recorded from above the resident cage. Testing ended after the first attack (defined as a bite) to prevent injury. If no attack occurred, the test ended at 10-minutes. The latency to attack was recorded as an indicator of aggression.

#### Statistical Analysis

All statistical analyses were performed using GraphPad Prism 9. For comparison of control to *Atrx*^NEXCre^ mice, a Students t-test was performed on time spent freezing for conditional fear memory; total distance travelled, average velocity, and distance travelled in center for the open field test; time spent in dark zone for the light/dark box paradigm; time spent grooming; %PPI; sociability index and number of tail rattling events for three-chamber test; agitation score; and latency to attack in the resident intruder paradigm. When variance was significant between samples, according to the F-statistic, a non-parametric Mann Whitney was performed. A two-way ANOVA was performed for distance over time analysis in the open field test, startle amplitude in the first portion of the PPI assessment, and for social preference in the three-chamber test. To evaluate sex differences, a discrimination index was calculated as follows: (*Atrx*^NEXCre^ value - average value for controls)/average value for controls. Discrimination indices for male and female mice were subjected to a Students t-test.

## Acknowledgments

We are grateful for access to the Neurobehavioural Core Facility at the Robarts Research Institute and for the support of facility manager Matthew Cowan.

## Data Availability

N/A

## Author Contributions

K.M.Q. contributed to conceptualization, design and execution of experiments, data interpretation, and writing of article. N.M.K. contributed to execution of experiments and data interpretation. N.G.B. contributed the conception, design, interpretation of data, and writing of article.

---

**Figure S1: Validation of ATRX deletion in *Atrx*^NEXCre^ female mice.**

**A)** ATRX immunofluorescence staining (red) of E13.5 control and *Atrx*^NEXCre^ brain coronal cryosections. DAPI counterstaining is shown in blue. Scale bar, 570 µm. **B)** Magnified view of ATRX (top, red) and βIII-tubulin (bottom, green) staining in the developing cortical plate. Scale bar, 144 µm. **C)** ATRX (red) and DAPI (blue) staining in the cortex and hippocampus of *Atrx*^NEXCre^ and control mice at P20. **D)** ATRX (red) and NeuN (green) staining in the cortex and hippocampus at P20. Cortex scale bar,100 µm and hippocampus scale bar, 400 µm. CA: cornu Ammonis, CP: cortical plate, Ctx: cortex, DG: dentate gyrus, GE: ganglionic eminence, LV: lateral ventricle, VZ: ventricular zone. eminence. Magnified images of cortical plate scale bar, 144 µm. VZ: ventricular zone, CP: cortical plate.

**Figure S2: Spatial learning deficits in *Atrx*^NexCre^ mice, without evidence of apathy.**

**A)** Latency to find the platform during the training phase of the Morris water maze test shows that *Atrx*^NEXCre^ male and female mice have impaired spatial learning (p<0.0001, F=3.630). **B)** *Atrx*^NexCre^ mice spend less time swimming during the trials, reflecting their floating behaviour (p<0.0001, F=19.50) (Control^male^ n=7, *Atrx*^NEXCre^ ^male^ n=7, Control^female^ n=7, *Atrx*^NEXCre^ ^female^ n=7). **C)** A 6-minute forced swim test does not reveal increased apathy in water for males (Control^male^ n=15, *Atrx*^NEXCre^ ^male^ n=13) and decreased apathy in females (p=0.0154, F=3.149, Control^female^ n=13, *Atrx*^NEXCre^ ^female^ n=12). Sex discrimination index indicates sex differences in the forced swim test (p=0.0026, F=1.010). Error bars represent +/-SEM. Three-way ANOVA or Students t-test performed when appropriate (* p<0.05, ** p<0.001, ***p<0.0001).

